# Volumetric development of hippocampal subfields and hippocampal white matter connectivity: Relationship with episodic memory

**DOI:** 10.1101/2022.01.31.478590

**Authors:** Anthony Steven Dick, Yvonne Ralph, Kristafor Farrant, Bethany Reeb-Sutherland, Shannon Pruden, Aaron T. Mattfeld

## Abstract

The hippocampus is a complex structure composed of several distinct subfields and has been at the center of scientific study examining the neural foundations of episodic memory. To date, there is little consensus regarding the structural development of the hippocampus and its subfields assessed both volumetrically and through anatomical connectivity, nor how the development of the related substructures influences episodic memory. In the current cross-sectional study, using a large sample of 830, 3- to 21-year-olds from a unique and publicly available dataset we examined the following questions: 1) Is there elevated grey matter volume of the hippocampus and its respective subfields in late compared to early development? 2) How does hippocampal volume compare with the rest of the cerebral cortex at different developmental stages? and 3) What is the relation between hippocampal volume, and the connectivity of the hippocampus with cortex as measured by diffusion-weighted imaging, with episodic memory performance? We found hippocampal subfield volumes exhibited a non-linear relation with age. Specifically, hippocampal subfield volumes showed a lag in volumetric change with age when compared to adjacent cortical regions (e.g., entorhinal cortex). We also observed a significant reduction in cortical volume across older cohorts, while hippocampal volume showed the opposite pattern. In addition to age-related differences in grey matter volume, several distinct subfields are significantly related to episodic memory. We did not, however, find any associations with episodic memory performance and connectivity through the uncinate fasciculus, fornix, or cingulum. The results are discussed in the context of current research and theories of hippocampal development and its relation to episodic memory.

## Introduction

The hippocampal formation and adjacent cortices that make up the medial temporal lobe (MTL) have long been known to play an important role in learning and memory (Scoville & Milner, 1957). Development of the MTL system, its maturation with respect to overall cerebral cortical development, and related functional consequences are central motivating questions. Here we attempt to 1) establish whether MTL development, with a focus on the hippocampus and respective subfields – both structurally and through white matter connectivity – continues beyond early childhood; 2) evaluate the relation between age related MTL volumetric differences with respect to the rest of the cerebral cortex; and 3) lastly assess how these age-related differences in volume and white matter connectivity relate to performance on a standardized memory test that assesses mechanisms important for episodic memory.

The MTL is a heterogeneous set of structures with unique functional attributes related to learning and memory comprised of the hippocampal formation and adjacent entorhinal, perirhinal, and parahippocampal cortices (Lech & Suchan, 2013; Squire, Stark, & Clark, 2004). The hippocampal formation can be further subdivided into distinct subfields defined by cytoarchitecture: cornu ammonis (CA) fields 1-3, dentate gyrus (DG), and subiculum (Winterburn et al., 2013). The unique functional contributions of the adjacent MTL cortices arise as a result of segregation of information, with spatial information predominately represented by the parahippocampal and medial entorhinal cortices and object information largely represented by the perirhinal and lateral entorhinal cortices (I. Lee, Hunsaker, & Kesner, 2005; Ranganath, 2010). The two streams of information subsequently converge on neurons in the hippocampal formation.

Distinct cytoarchitecture of the hippocampal subfields further confers functional attributes to the MTL system (Marr, 1971). The DG is comprised of a high volume of largely quiescent cells. The large number of neurons, combined with a low firing rate, provide the ideal substrate for performing a computation known as “pattern separation,” which is proposed to orthogonalize or make overlapping representations distinct (E. Rolls, 2013; E. T.Rolls, 1996). Neurophysiological evidence of pattern separation has been identified in both rodent and human studies (Bakker, Kirwan, Miller, & Stark, 2008; Leutgeb, Leutgeb, Moser, & Moser, 2007). Neurogenesis, which occurs in only a handful of places in the adult brain, including the DG, is also thought to play an important role in pattern separation (Pereira et al., 2007). In contrast, the most notable cytoarchitectural feature of the CA3 subfield is its recurrent collaterals (Bains, Longacher, & Staley, 1999). The structure of the CA3 makes this subfield adept at a computation called “pattern completion” or the reinstatement of a prior representation from partial input (E. T. Rolls, 1996). Lastly, the CA1 subfield receives notable input from both the CA3 and entorhinal cortex (Brun et al., 2008), distinguishing this subfield as a comparator of current information (incoming from the entorhinal cortex) with previously encoded information (retrieved through the CA3 process of pattern completion; Lisman & Grace, 2005). Together these regions and their unique functional attributes support the ability to discriminate overlapping memories, generalize across similar episodes, and lay down new memories.

### Differential Development of the Hippocampus, Its Subfields, and Its Connectivity

In addition to investigating the functional attributions to subdivisions of the hippocampus, histological studies in non-human primates (Lavenex & Lavenex, 2013; Lavenex, Lavenex, & Amaral, 2007) combined with cross-sectional and longitudinal human MRI studies (see Jones & McHugh, 2011 for a review) have observed distinct developmental trajectories of the cortex and hippocampal formation as well as the different hippocampal subfields. Several important questions have been raised in the literature, and some have been answered to varying degrees. The present study focuses on a number of questions that remain unanswered, mainly due to the presence of low statistical power in the extant literature, or to the investigation of restricted age ranges (Joshua K Lee, Johnson, & Ghetti, 2017; Uematsu et al., 2012). These open questions are: 1) does the structure of the hippocampus continue to develop substantially beyond early childhood? (Joshua K Lee, Ekstrom, & Ghetti, 2014); 2) how does the structure of the hippocampus develop in relation to the rest of the cerebral cortex; 3) does the white matter connectivity of the hippocampus develop substantially beyond early childhood; and 4) how do these factors relate to the development of episodic memory at the behavioral level?

### Hippocampal structural development

There are a number of studies that have examined the hippocampus as a whole structure, yet these studies have failed to come to a consensus on whether the hippocampus continues to show age-related changes throughout development (Goddings et al., 2014; Joshua K Lee et al., 2015; Østby et al., 2009; Riggins et al., 2018; Tamnes et al., 2013; Uematsu et al., 2012; Yurgelun-Todd, Killgore, & Cintron, 2003). Some studies have found that development of the hippocampus peaks earlier in childhood or have otherwise failed to find age-related differences, even when substantial age ranges are investigated (Barnea-Goraly et al., 2014; Giedd et al., 1996; Gogtay et al., 2006; Joshua K Lee et al., 2015; Riggins, Blankenship, Mulligan, Rice, & Redcay, 2015; Uematsu et al., 2012). For example, in a longitudinal study of 31 children with repeated scans, Gogtay and colleagues (2006) reported that the total volume of the hippocampus remained unchanged bilaterally from age 4 to 25 years. However, other studies investigating similarly broad age ranges have found substantial age-related changes (e.g., Østby et al., (2009). For example, in a large cross-sectional study of 171 participants aged 8 to 30 years, Østby and colleagues (2009) reported non-linear decreases in grey matter in cerebral cortex, with concomitant increases in overall volume in amygdala and hippocampus, peaking during adolescence. This non-linear developmental trajectory was later replicated in a large cross-sectional morphometric MRI study of the amygdala and hippocampus from 1 month to 25 years of age (Uematsu et al., 2012). Thus, there remain considerable discrepancies in the literature regarding how the structure of the hippocampus changes with development.

More recently, a growing body of literature has investigated structural development of the hippocampus with increased attention to the possibility of differential development of hippocampal subfields. Riggins and colleagues (2018) examined younger children between 4 and 9 years, and found age effects only in CA1. Lee and colleagues (2014) demonstrated age-related increases, in a sample of 39 eight- to fourteen-year-olds, in CA3 and DG between middle childhood and early adolescence, consistent with previous histologic studies (c.f. Insausti, Cebada-Sanchez, & Marcos, 2010). Similarly, Krogsrud and colleagues (2014) demonstrated in 244 healthy participants, aged between 4 and 22 years, that there is a rapid development of most of the hippocampal subfields until around 13 to 15 years. In a recent update to their earlier work, Canada and colleagues (Canada, Botdorf, & Riggins, 2020; Canada, Hancock, & Riggins, 2021) examined the development of hippocampal head, body, and tail in a longitudinal design, and reported increases in volume during early-to-middle childhood (up to age 9 years), although the nature of the change was more linear in the body and tail. Together, these results are consistent with some of the results we discussed above suggesting protracted development of the hippocampus and its subfields into middle childhood and early adolescence.

An important question, which has been addressed in some prior work, is how the development of hippocampal subfields might relate to episodic memory development (see Ghetti & Bunge, 2012 for a review). In recent work, DeMaster and colleagues (2013) investigated this issue in a sample of 37 eight- to eleven-year-olds and 25 adults. They found a different pattern in the relation between memory performance in adults and children, with memory performance more associated with smaller right hippocampal head, and larger hippocampal body, in adults, but not children. In children, but not adults, this association was found for the left hippocampal tail. More recently, Riggins et al. (2018) examined the relation between age-related hippocampal structural changes and memory development in younger children. Overall, there was a negative relation between memory performance and CA1 volume in the body, and positive relation between memory performance and CA2-4/DG volume in the body. These differential relations highlight the need, as Riggins and colleagues suggest, to examine the subregions of the hippocampus in relation to performance. Examination of the whole hippocampus may lead to an underestimation of the relation between memory performance and the distinct circuits subserved by these subregions.

Less attention has been paid to development of the fiber pathways connecting the hippocampus and its subfields to the rest of the cortex. Three major fiber pathways establish connectivity of the hippocampus with other subcortical structures and of the cortex, and these may be important for episodic memory development. These are the fornix, which connects hippocampus to the thalamus (Douet & Chang, 2015), hippocampal cingulum, which connects the hippocampus and parahippocampal cortex with the cingulate cortex (Jones et al. 2013), and uncinate fasciculus, which connects the hippocampus to the anterior temporal and inferior frontal lobes (Dick, Bernal, & Tremblay, 2014; Dick & Tremblay, 2012; Olson, Von Der Heide, Alm, & Vyas, 2015). Lebel and colleagues (2012) have reviewed studies of age-related structural changes in these fiber pathways and show the fornix, as measured by fractional anisotropy (FA), is one of the earliest fiber pathways to reach an asymptote in terms of age-related change, peaking at around 19 years. The uncinate fasciculus, on the other hand, peaked around 30 years, and the cingulum continued to evidence age-related changes until about 42 years. None of these studies though investigated how these changes were associated with concomitant changes in hippocampal structure, and only limited work has been conducted investigating the relation between development in these pathways and memory development (Mabbott, Rovet, Noseworthy, Smith, & Rockel, 2009; Ngo et al., 2017; Wendelken et al., 2014).

In summary, despite a number of well-designed studies, how the volume of the hippocampus changes over a considerable age range, and within distinct subfields, and how this plays a role in episodic memory, remains underspecified. Furthermore, the relation between white matter connectivity of the hippocampus over development, and its association with episodic memory development, remains poorly understood. Thus, to address the extant questions, the present study uses a very large dataset, across a large age range, to examine the developmental trajectory of the hippocampus as a whole, hippocampal subfields, and hippocampal connectivity, and how these indices relate to episodic memory performance as a function of age.

## Method and Design

### Participants

The current cross-sectional study was conducted using MRI data from the publicly available database, Pediatric Imaging, Neurocognition, and Genetics (PING) (http://pingstudy.ucsd.edu). Data were provided on an external hard-drive that included a virtual machine and Linux mountable partition. Data from the Linux partition were uploaded to our local high-performance computing cluster. The PING database consists of 1493 typically-developing participants between the ages of 3.0- and 21.0-years-old (713 female). Participants were scheduled for two separate visits. The first included neurophysiological assessments and the second included MRI data acquisition. Of these 1493 participants, our analysis was conducted on a total of 830 participants (380 female), with an age range of 3.0 to 21.0 years (*M* = 12.6; *SD* = 4.9). Age was recorded at the time of the MRI scan. The discrepancy between the number of participants available and the number of participants in our analysis is due to the fact that 830 participants had an available MRI data that had been processed through FreeSurfer so that we could conduct the hippocampal segmentation (see below).

As part of the PING study, participants were administered multiple neuropsychological tests, including social and emotional assessments from the Phoenix toolkit (https://www.phenxtoolkit.org) and cognitive assessments from the NIH Toolbox (http://www.nihtoolbox.org). In this study, we focused on the NIH Toolbox Picture Sequence Memory Task (PSMT), which is based on the Imitation Based Assessment of Learning (IBAL), a measure of episodic memory.

### Image acquisition and Data Analysis

Image acquisition and image data post-processing were conducted by the PING team as described in Jernigan et al., (2016). We provide only cursory details here. Across 10 sites and 12 scanners, a standardized, multiple modality high-resolution MRI protocol was implemented using the following scanner manufactures: GE Discovery MR 750, GE Signa HDX, Siemens Trio TIM, and Phillips Medical Systems Achieva. T1-weighted structural and diffusion weighted imaging (DWI) data were used for the present study. The scanning protocols are summarized as follows: Siemens (T1-weighted: TR=2170ms, TE=4.33ms, flip angle=7°, matrix size=256×256mm, voxel size: 1×1×1.2, acquisition time = 8:06 minutes; Diffusion-weighted: Isotropic (2.5×2.5×2.5mm), single shot echo-planar sequence protocol: slices=68, slice thickness=2.5mm, FOV=240×240×170mm, TR=9000ms, TE=91ms, directions=30, b=1000 s/mm^2^, acquisition time = 10:00 minutes, acquisition matrix = 240×240×170); Phillips (T1-weighted: TR=6.8ms, TE=3.1ms, flip angle=8°, matrix size=256×240mm, voxel size: 1×1×1.2, acquisition time: 9:19 minutes; Diffusion-weighted: Isotropic (2.5×2.5×2.5mm), single shot echo-planar sequence protocol: slices=60, slice thickness=2.5mm, FOV=240×240×150mm, TR=9000ms, TE=91ms, directions=30, b=1000 s/mm^2^, acquisition time = 5:15 minutes, acquisition matrix = 96 x95); GE (T1-weighted: TR=8.1ms, TE=3.5ms, flip angle=8°, matrix size=256×192mm, voxel size: 1 x1×1.2, acquisition time: 8.05 minutes. Diffusion-weighted: Isotropic (2.5×2.5×2.5mm), single shot echo-planar sequence protocol: slices=60, slice thickness=2.5mm, FOV=240×240×150mm, TR=9000ms, TE=91ms, directions=30, b=1000 s/mm^2^) acquisition time: 5:15 minutes, acquisition matrix = 96 x95).

### Post-processing and quality control of MRI data

All post-processing of the volumetric and diffusion data was conducted by the PING study team and is described in Jernigan et al. (2016) with the exception that the hippocampal segmentation was conducted by us (described below). A standard PING scan session included: 1) a 3D T1-weighted inversion prepared RF-spoiled gradient echo scan using prospective motion correction (PROMO; White et al., 2010) for cortical and subcortical segmentation; 2) a 3D T2-weighted variable flip angle fast spin echo scan, which also used PROMO for detection and quantification of white matter lesions and segmentation of CSF; and 3) a high angular resolution diffusion imaging (HARDI) scan, with integrated B__0__ distortion correction (DISCO), for segmentation of white matter tracts and the measurement of diffusion parameters.

Raw image quality control was conducted for each scan session of the PING project. Images were automatically checked for completeness and protocol compliance, and were reviewed for image quality by technicians. All images were screened for motion artifacts, excessive distortion operator error, or scanner malfunction. Images were rated with either, good, average, and bad. T1-weighted images were examined slice by slice for excessive motion. Each volume was rated as either as acceptable or recommended for rescan.

Because we wanted to take advantage of this extensive post-processing pipeline, we began our investigation with T1-weighted scans that had been processed through FreeSurfer 5.3 and which were part of the public release. FreeSurfer: a) segments the white and grey matter of anatomical volumes; b) inflates the cortical surfaces separately for each hemisphere (Dale, Fischl, & Sereno, 1999; Fischl, Sereno, Tootell, & Dale, 1999); and c) segments the subcortical structures, including the hippocampus. We further applied the hippocampal segmentation in FreeSurfer v6.0, as described below.

### Segmentation of the hippocampus

For the structural analysis of hippocampal volume, we re-ran the reconstruction of the v5.3 public release data using FreeSurfer v6.0, including the hippocampal segmentation step to segment the hippocampus into subfields. This step follows the routine outlined in Inglesias et al. (2015) and was invoked by applying the --hippocampal-subfields-T1 flag to the standard FreeSurfer reconstruction. This reconstruction was applied to all 830 participants who had available FreeSurfer v5.3 reconstructions as part of the public release. The entire recon-all procedure was applied in v6.0, beginning with the T1.mgz that was provided with the release (thus, it was already subjected to the quality control steps through PING). The hippocampal subfield segmentation in v6.0 employs a number of substantial improvements over v5.3, which was the first version of FreeSurfer to include a hippocampal subfield segmentation option. These improvements, which specify the training atlas used for automated segmentation, include use of ultra-high resolution (0.1 mm isotropic) *ex vivo* MRI data collected at 7T in 15 brains (employing 0.13 mm isotropic resolution, allowing delineation of a molecular layer), delineation of a large number of structures (i.e., 15), and modeling of surrounding structures to constrain labeling within the hippocampus. An unbiased segmentation of each brain in our sample was computed based on this training atlas using a Bayesian inference algorithm (Iglesias et al., 2015). The procedure was validated on three publicly available data sets and has been found to have good agreement to manual labeling of *ex vivo* scans (Iglesias et al., 2015). Using this automated tool, we segmented the hippocampal formation into the following sub-divisions, with DG and CA3 combined: CA1, DG/CA3, subiculum, and entorhinal cortex (not part of hippocampus proper) in both hemispheres.

### Diffusion-weighted imaging data post-processing

Diffusion data were post-processed by the PING team with the following steps: spatial and intensity distortions were reduced by application of non-linear registration to a reverse phase-encoded scan (Holland, Kuperman, & Dale, 2010). Eddy current distortions were corrected with a nonlinear estimation procedure (Hagler et al., 2009). See Jernigan et al (2016) for additional details.

### Diffusion tensor imaging analysis

All of the fiber pathways in the current study were traced by the PING team. To trace these pathways, the PING team employed the automated labelling process developed by Hagler et al., (2009). This method uses a probabilistic atlas containing 23 fiber tracts which were constructed by manually identifying fiber tracts in 21 healthy controls and 21 patients with temporal lobe epilepsy (for additional details please see (Hagler et al., 2009). The tracking methods include the principal diffusion orientation, fractional anisotropy (FA), mean, longitudinal, and transverse diffusivity (MD, LD, and TD). To label long-range white matter tracts, AtlasTrack was used to automatically label fiber pathways based on a probabilistic atlas of fiber tract locations and orientations (Hagler et al., 2009).

### Behavioral methods

To examine the relation between volumetric and diffusion weighted images and episodic memory performance, we used the NIH Toolbox Picture Sequence Memory Test (PSMT), conducted as part of the NIH Toolbox Cognition battery (http://nihtoolbox.com), and gathered as part of the PING data collection efforts. This task is similar to the Imitation Based Assessment of Learning (IBAL) developed by Bauer and colleagues (2013). In this task, images of pictured objects and activities are presented on an iPad in an arbitrary order. The participant’s task is to view the presentation of the images, and then to reproduce the ordering of the images after they leave the screen. The sequence length is varied to manipulate task difficulty, from 6 to 15 pictures (Bauer et al., 2013). The participant’s score is derived from the cumulative number of adjacent pictures remembered correctly over three learning trials. The task has good test-retest reliability (ICC = .76) and convergent validity with other measures of episodic memory (Bauer et al., 2013).

### Statistical analysis

Three separate analyses were performed. The first set of analyses examined age-related differences in the volume of the whole hippocampus and entorhinal cortex, and the hippocampus segmented into three subregions (CA1, DG/CA3, and subiculum). The second set of analyses examined age-related differences in FA of the three fiber pathways of interest: the fornix, the uncinate fasciculus, and the hippocampal cingulum. The third set of analyses examined the relation of these morphologic and diffusion metrics to performance on the NIH Toolbox Picture Sequence Memory Task.

#### ComBat Correction for Cross-Site Differences

Before conducting the analysis, we also corrected the data (both morphologic and diffusion) for cross-site differences in scanner type and acquisition protocols. To make this correction, we employed the ComBat method developed by Johnson and colleagues (2007) and applied to imaging data by Fortin and colleagues (2017). This method uses an empirical Bayes framework to calculate a location and scale adjustment to the data. It estimates an empirical statistical distribution of the location and scale parameters under the assumption that all voxels share a common distribution. Within a regression framework, an adjusted value is computed for each data point. The ComBat adjusted data were used for all analyses going forward.

#### Analysis 1: Age-related differences in volume of hippocampus and entorhinal cortex

To analyze age-related differences in hippocampal, entorhinal cortex, and hippocampal subregion volume (for both hemispheres), we followed the procedure from Østby and colleagues (2009) and corrected for residual intracranial volume (ICV). This is done because brain structures tend to scale with head size, and we are interested in size differences of the structures of interest that are not confounded with head-size differences. The estimate of ICV was taken from the FreeSurfer output, and was calculated by using atlas normalization as a proxy for ICV (see Buckner et al., 2004).

We used the residual method to correct for differences in head size (Sanfilipo, Benedict, Zivadinov, & Bakshi, 2004). Each volume of interest (dependent variable) was subjected to a regression, with ICV predicting the dependent variable. The residuals of this regression were standardized, and passed to the next analysis (i.e., the new dependent variable is a scaled ICV-corrected volume estimate for each structure of interest). A generalized additive model (GAM; R 3.4.3; package *gam*; R Core Team, 2017) was fit to each variable of interest, with age at exam as the predictor. Integrated smoothness estimation was applied (of the form *gam*(*dv*~*s*(*predictor*)). We computed Akaike Information Criterion (AIC) values for each model. In addition, we computed the first derivative of the data to identify the “peak” of the curve, in cases where the function was non-linear.

We also conducted an analysis of the hippocampus proper in relation to development of the cortex. To do so we computed the ratio of hippocampal volume to volume of the cortex (hippocampal volume/total cortical volume). Cortical volume was derived from FreeSurfer by taking the volume of tissue inside the pial brain surface minus the volume of tissue inside the white matter surface, and minus the volume of non-cortical grey matter (e.g., the hippocampus). This variable was scaled and subjected to the same GAM analysis as above.

#### Analysis 2: Age-related differences in FA of the fornix, uncinate fasciculus, and hippocampal cingulum

We used a similar approach as we described above, except that whole brain FA was residualized in place of ICV. Thus, in each case, for each hemisphere, we specified age as the predictor in the model, and residualized scaled FA of each pathway as the outcome variable. The first derivative was similarly computed for each of the pathways, for each hemisphere.

#### Analysis 3: Relation of morphologic and diffusion metrics to performance on the NIH Toolbox Picture Sequence Memory Task

To explore the relation between morphologic and diffusion metrics and episodic memory, we conducted linear mixed effects models (R package *lme*) using each hippocampal volume measure, FA in each pathway of interest, and entorhinal cortex volume, for each hemisphere. Age, gender, ethnicity, parent highest level of education, household income, and device serial number (to control for scanner type) were entered as covariates. In addition, ICV (for the morphologic variables) and whole brain FA (for the diffusion variables) were entered as covariates. Finally, subject ID was entered as a random effect. We also ran these analyses using hippocampal-cortical ratio as a predictor.

## Results

### Results of Analysis 1: Age-related differences in volume of hippocampus and entorhinal cortex

The results of Analysis 1 are presented in Figure 1 and Table 2. These results reflect the volume of the regions of interest after controlling for ICV. The analysis showed that for the whole hippocampus, in both hemispheres, there is a similar non-linear relation such that the volume of the hippocampus is elevated with increasing age up until 15 years-old and then plateaus. Similar trajectories were demonstrated for CA1, DG/CA3, and subiculum, with volumes peaking in the 13- to 15-year age range. The entorhinal cortex did not show such a trajectory, and instead evidenced a flat trend, indicating it did not show age-related differences that were dissociable from the changes seen in the rest of the cortex. Results of the analysis of hippocampal-cortex volume ratio are presented in Figure 2 and Table 2. These are plotted against the age-related differences seen at the whole-brain level (grey line). For the ratio, the age-related difference trajectory was linear, with no evidence of an asymptote. Thus, the ratio of the hippocampal volume relative to the cortex reflects a continued increase with age.

**Table 2.** Model fit for the linear and quadratic models for each region and pathway.

| <u>Region</u> | Hemisp<br>here | Linear Model AIC | Quadratic Model AIC |
| --- | --- | --- | --- |
| Hippocampus | Left | 2325.7 *** | **2313.7 \*\*\*** |
|  | Right | 2328.8 *** | **2320.7 \*\*\*** |
| Entorhinal Cortex | Left | 2349.7 *n.s.* | 2349.7 *n.s.* |
|  | Right | 2349.8 *n.s.* | 2348.9 *n.s.* |
| CA1 | Left | 2333.6 *** | **2319.8 \*\*\*** |
|  | Right | 2324.8 *** | **2319.7 \*\*\*** |
| Dentate gyrus/CA3 | Left | 2314.3 *** | **2298.4 \*\*\*** |
|  | Right | 2313.5 *** | **2302.9 \*\*\*** |
| Subiculum | Left | 2342.8 ** | **2340.4 \*** |
|  | Right | 2342.8 ** | **2338.2 \*\*** |
| Hippocampal/Cortical Ratio | Left | 2045.6 *** | 2045.6 *** |
|  | Right | 2049.9 *** | 2049.1 *** |
| <u>Pathway</u> |  |  |  |
| Hippocampal cingulum | Left | 2338.8 ** | **2331.1 \*\*** |
|  | Right | 2334.8 ** | **2331.5 \*\*** |
| Fornix | Left | 2333.0 ** | **2332.6 \*\*** |
|  | Right | 2318.9 *** | 2318.9 *** |
| Uncinate fasciculus | Left | 2344.3 *n.s.* | 2340.3 *n.s.* |
|  | Right | 2346.8 * | 2344.7 *n.s.* |

*Note.* \* $p < .05$ ; \*\* $p < .01$ ; \*\*\* $p < .001$ ; *n.s.* = non-significant. Bold denotes the smaller Akaike Information Criterion (AIC), indicating better model fit.

**Figure 1.**
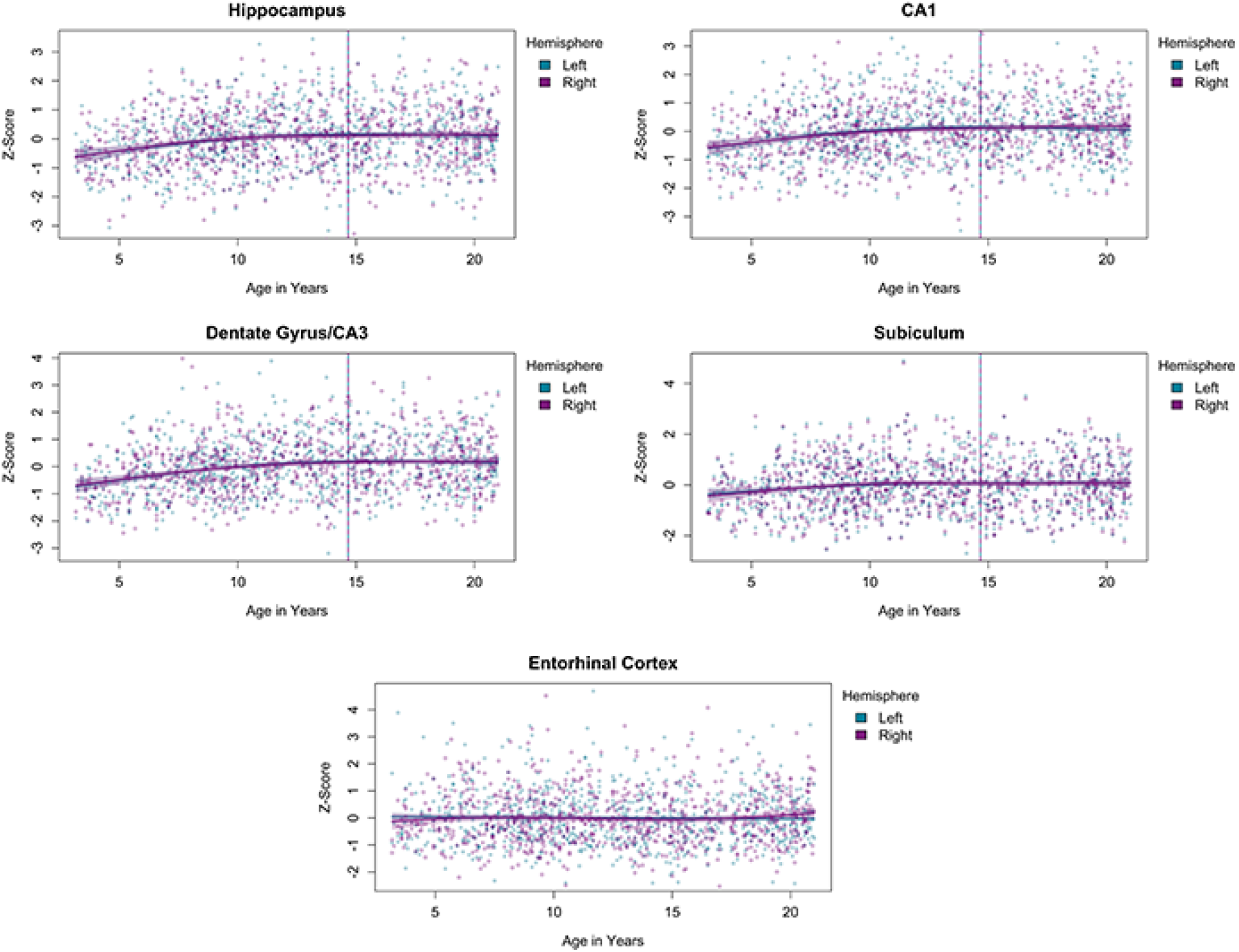
Age-related differences in bilateral hippocampus, each hippocampal subfield (CA1, DG/CA3, Subiculum) and entorhinal cortex. Results reflect the volumes of the regions of interest after controlling for whole-head intracranial volume (ICV). The vertical line indicates the calculated first derivative for each curve.

**Figure 2.**
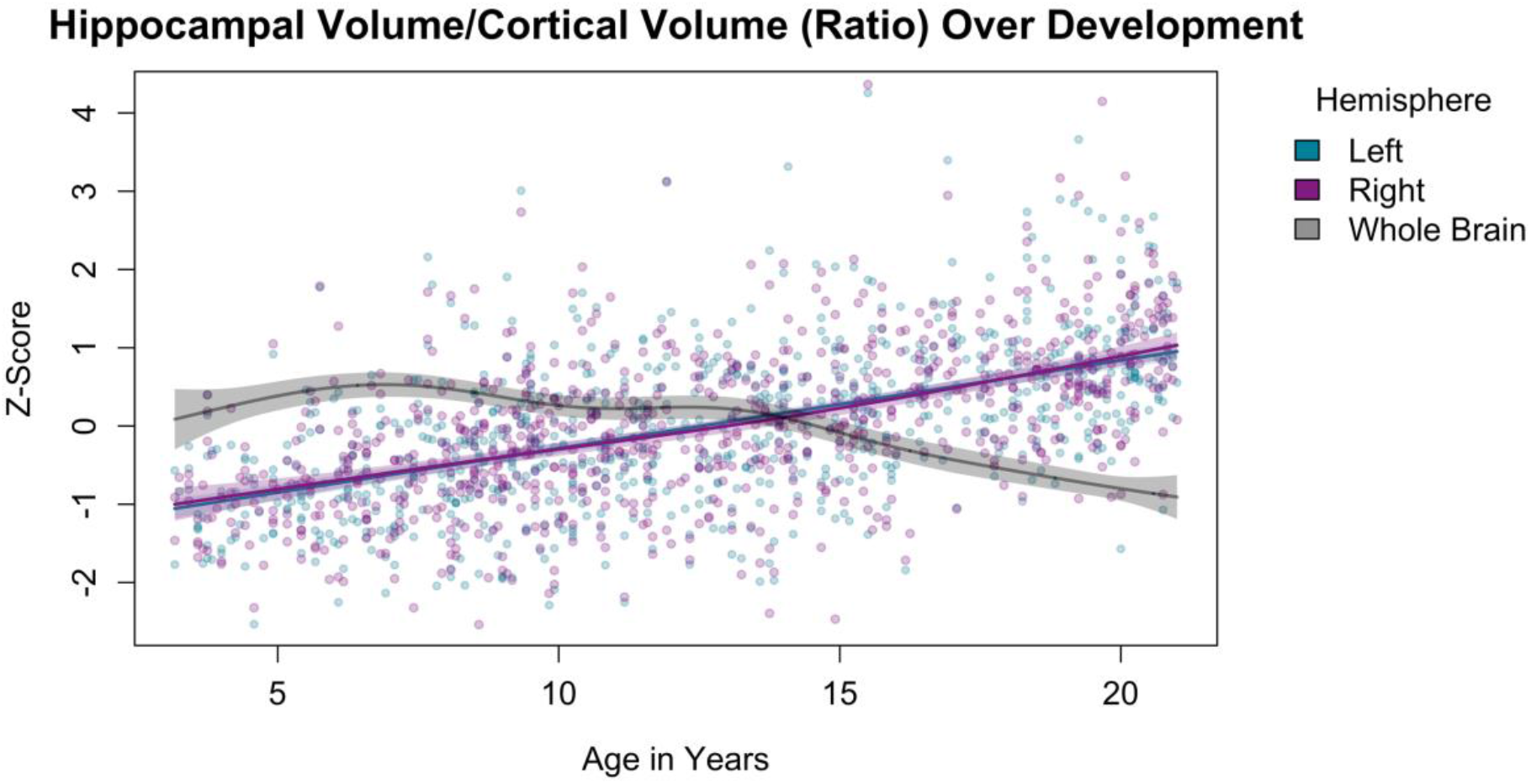
Age-related differences for the bilateral hippocampal-cortical ratio. Age-related differences in whole-brain volume are plotted in grey.

### Results of Analysis 2: Age-related differences in FA of the hippocampal cingulum, fornix, and uncinate fasciculus

The results of Analysis 2 are presented in Figure 3 and Table 2. These results reflect the FA of the pathways of interest after controlling for whole-brain FA. The analysis showed (bilaterally) a positive slope for the cingulum, a negative slope for the fornix, and no discernable age-related differences for the uncinate fasciculus. However, even for the cingulum and fornix, the changes over this age range were small, and no asymptote was apparent.

**Figure 3.**
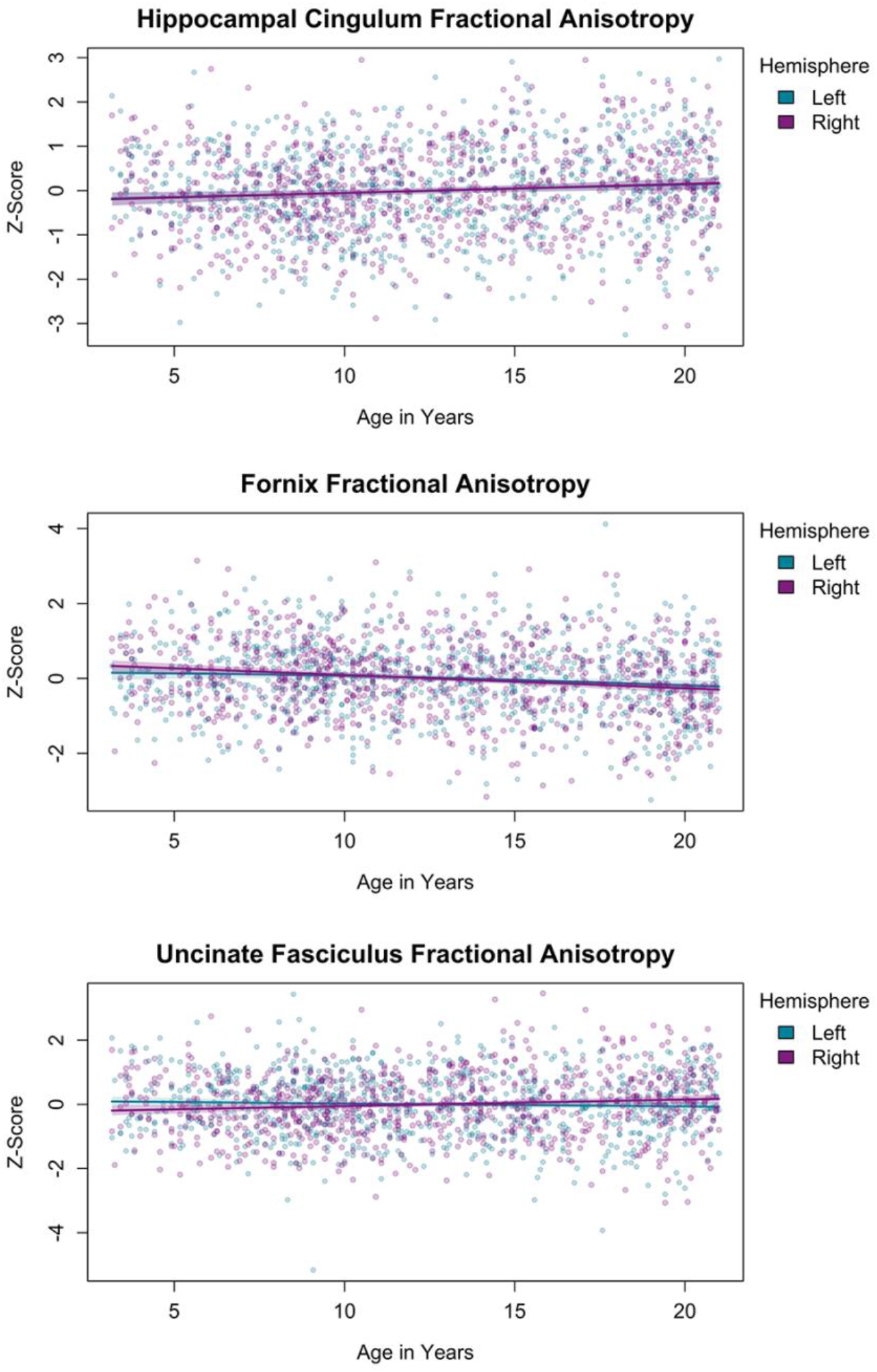
Age-related differences in fractional anisotropy (FA) of the hippocampal cingulum, fornix, and uncinate fasciculus, controlling for whole-brain FA.

### Results of Analysis 3: Relation of morphologic and diffusion metrics to performance on the NIH Toolbox Picture Sequence Memory Task

The results of Analysis 3 are presented in Figure 4 and Table 3. The volume of bilateral hippocampus, bilateral DG, left CA1, and right DG/CA3 were positively associated with performance on the PSMT, our episodic memory task. The ratio of hippocampal to cortical volume (bilaterally) was also positively associated with performance on the PSMT. We did not find any relations between FA of any of the fiber pathways and episodic memory performance.

**Table 3.**
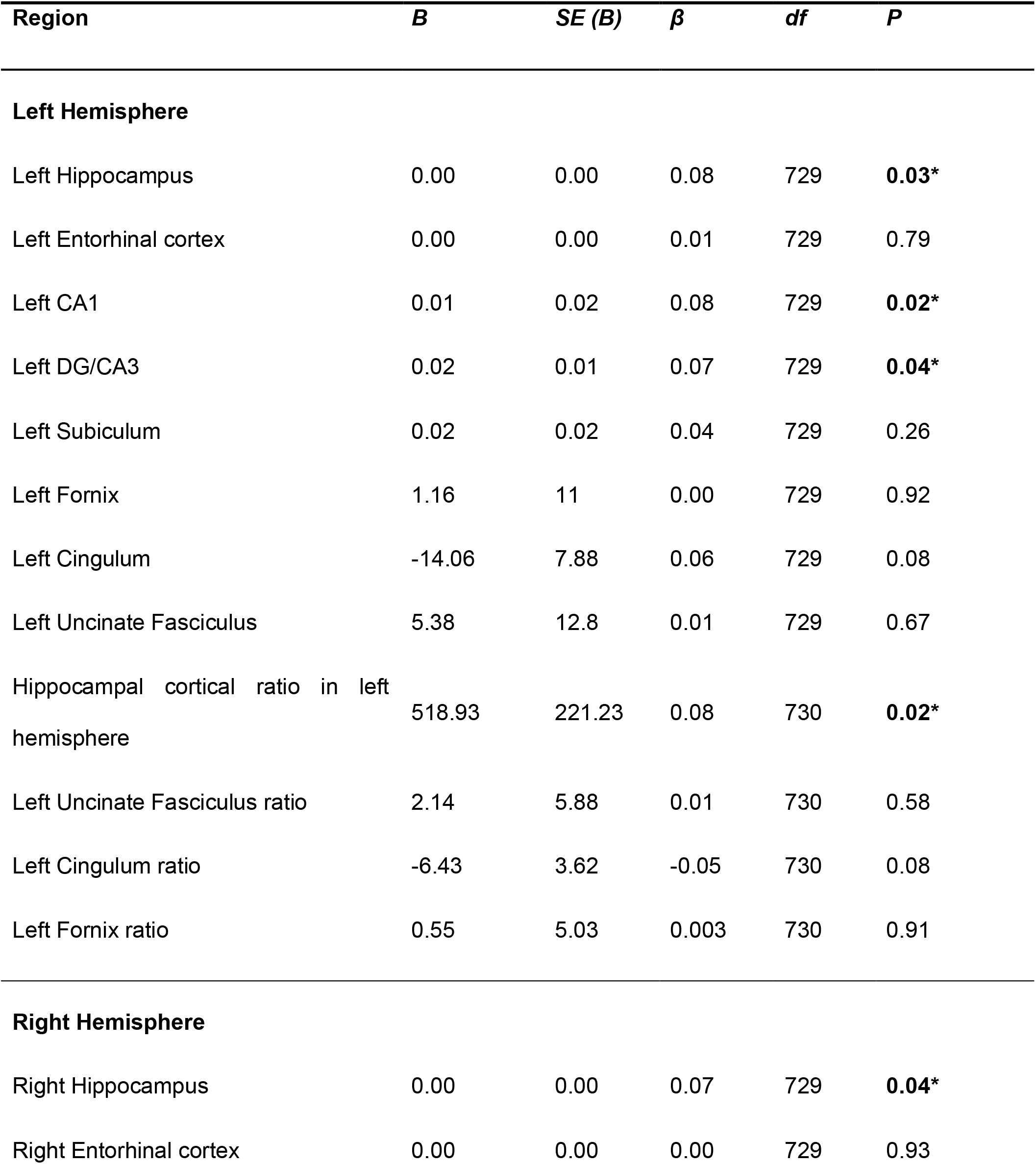
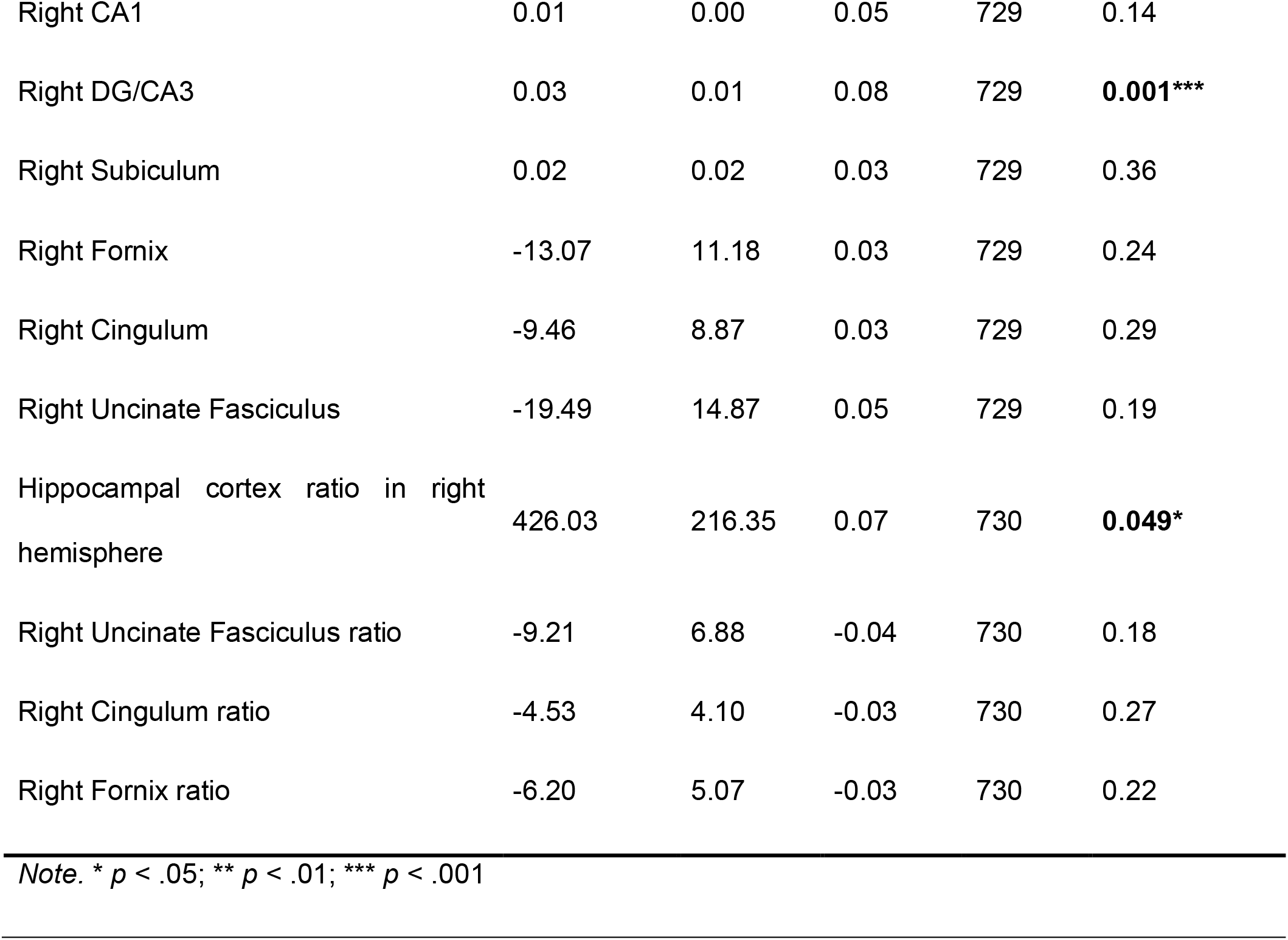
Summary of regression results for regions and pathways of interest predicting episodic memory performance.

| Region | *B* | *SE (B)* | $\beta$ | *df* | *P* |
| --- | --- | --- | --- | --- | --- |
| **Left Hemisphere** |  |  |  |  |  |
| Left Hippocampus | 0.00 | 0.00 | 0.08 | 729 | **0.03\*** |
| Left Entorhinal cortex | 0.00 | 0.00 | 0.01 | 729 | 0.79 |
| Left CA1 | 0.01 | 0.02 | 0.08 | 729 | **0.02\*** |
| Left DG/CA3 | 0.02 | 0.01 | 0.07 | 729 | **0.04\*** |
| Left Subiculum | 0.02 | 0.02 | 0.04 | 729 | 0.26 |
| Left Fornix | 1.16 | 11 | 0.00 | 729 | 0.92 |
| Left Cingulum | -14.06 | 7.88 | 0.06 | 729 | 0.08 |
| Left Uncinate Fasciculus | 5.38 | 12.8 | 0.01 | 729 | 0.67 |
| Hippocampal cortical ratio in left hemisphere | 518.93 | 221.23 | 0.08 | 730 | **0.02\*** |
| Left Uncinate Fasciculus ratio | 2.14 | 5.88 | 0.01 | 730 | 0.58 |
| Left Cingulum ratio | -6.43 | 3.62 | -0.05 | 730 | 0.08 |
| Left Fornix ratio | 0.55 | 5.03 | 0.003 | 730 | 0.91 |
| **Right Hemisphere** |  |  |  |  |  |
| Right Hippocampus | 0.00 | 0.00 | 0.07 | 729 | **0.04\*** |
| Right Entorhinal cortex | 0.00 | 0.00 | 0.00 | 729 | 0.93 |
| Right CA1 | 0.01 | 0.00 | 0.05 | 729 | 0.14 |
| Right DG/CA3 | 0.03 | 0.01 | 0.08 | 729 | **0.001\*\*\*** |
| Right Subiculum | 0.02 | 0.02 | 0.03 | 729 | 0.36 |
| Right Fornix | -13.07 | 11.18 | 0.03 | 729 | 0.24 |
| Right Cingulum | -9.46 | 8.87 | 0.03 | 729 | 0.29 |
| Right Uncinate Fasciculus | -19.49 | 14.87 | 0.05 | 729 | 0.19 |
| Hippocampal cortex ratio in right hemisphere | 426.03 | 216.35 | 0.07 | 730 | **0.049\*** |
| Right Uncinate Fasciculus ratio | -9.21 | 6.88 | -0.04 | 730 | 0.18 |
| Right Cingulum ratio | -4.53 | 4.10 | -0.03 | 730 | 0.27 |
| Right Fornix ratio | -6.20 | 5.07 | -0.03 | 730 | 0.22 |

*Note.* \* $p < .05$ ; \*\* $p < .01$ ; \*\*\* $p < .001$

**Figure 4.**
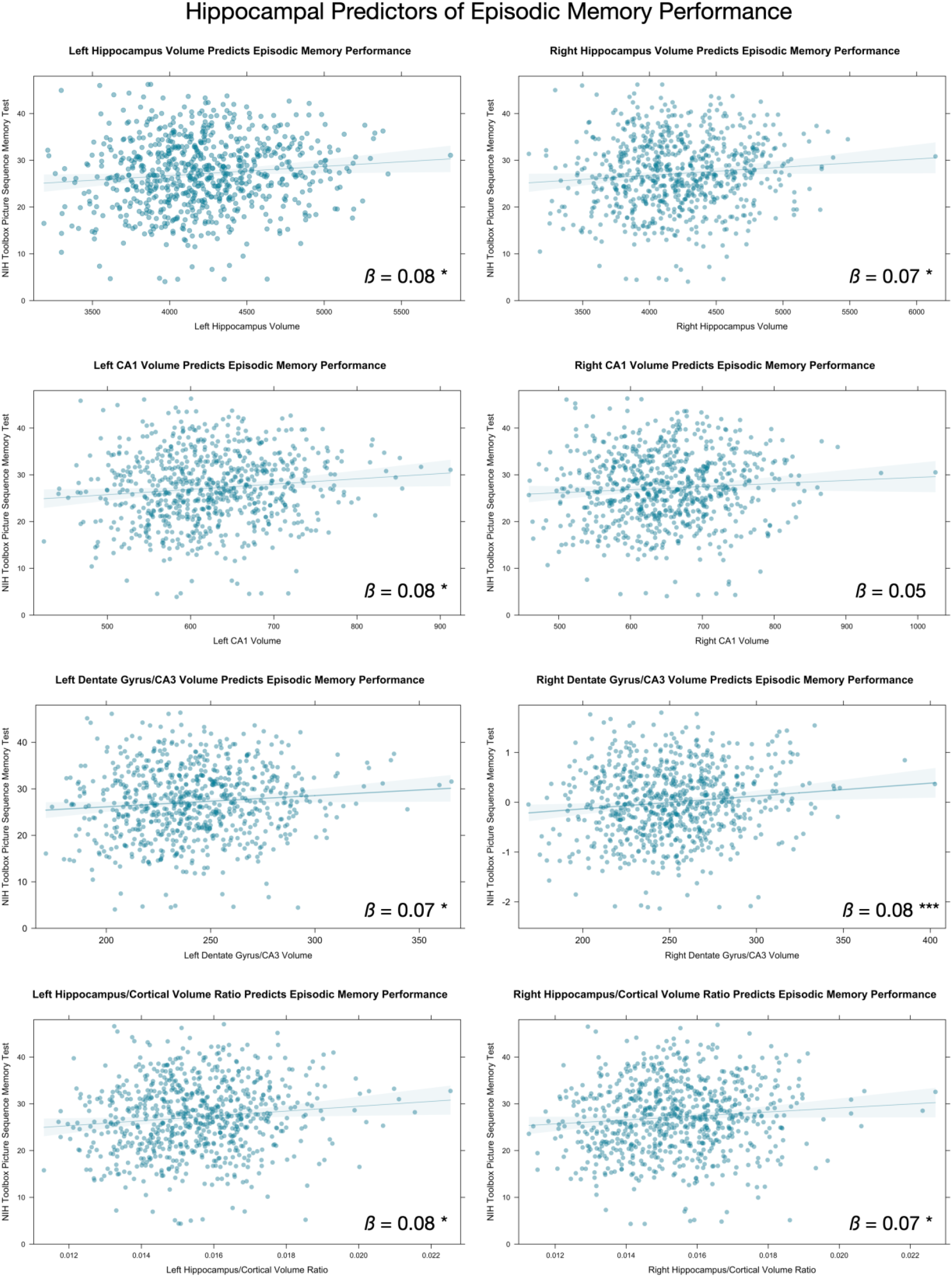
Associations between hippocampus and subregion volume, and episodic memory performance. Only significant associations from Table 3 are shown. * *p* < .05. *** *p* < .001.

## Discussion

The hippocampus is a complex structure composed of several distinct subfields and has been at the center of scientific study examining the neural foundations of episodic memory. In order to understand episodic memory, it is necessary to establish the boundaries that define typical development of the structures that support it, namely the hippocampus. Here, in a large sample of 830 three- to twenty-one-year-olds from a unique and publicly available dataset, we examined age-related differences in the volume of the hippocampus, its subfields, and in the diffusion properties of the fiber pathways connecting hippocampus to other cortical and subcortical structures. We found greater hippocampal and subfield volume exhibiting a non-linear relation with age peaking in the mid-teen years. A similar relation between age and volume in an adjacent cortical region, the entorhinal cortex, was not observed. We also found that while cortical volume decreased with age, hippocampus volume continued to increase. Age-related elevations in grey matter volume, across MTL structures was significantly positively related to episodic memory performance. Finally, we examined how MTL white matter pathways were associated with episodic memory, but did not find any evidence, in this dataset, of a significant relation between these pathways and age nor episodic memory performance. We discuss each of our findings in turn.

### 1. Hippocampal subfield and entorhinal cortex grey matter volume development

We first examined the basic volumetric changes associated with age in regions supporting episodic memory, including the entorhinal cortex, and the hippocampus and its subfields. We found non-linear developmental trajectories which continue to evidence greater volumes with age well into the mid-teen years for the whole hippocampus, as well as for the hippocampal subfields CA1 and DG/CA3. We did not see the same protracted increase in volume in either the subiculum or the entorhinal cortex, the latter the major cortical input to the hippocampus.

Whole-hippocampal volume exhibits a protracted developmental trajectory. Many studies have examined the structural development of the hippocampus (Goddings et al., 2014; Joshua K Lee et al., 2015; Østby et al., 2009; Tamnes et al., 2013; Uematsu et al., 2012; Yurgelun-Todd et al., 2003). Our findings – greater age-related hippocampal volumes well into mid-teen years – are consistent with prior work that similarly observed non-linear age-related increases in volume that peaked in adolescence (Østby et al., 2009; Uematsu et al., 2012). However, examining the hippocampus as a whole structure has yielded somewhat inconsistent results, especially in younger samples. For example, Gogtay and colleagues (2006) reported no age-related changes in overall hippocampal volume between 4 and 25 years, while Tamnes and colleagues (2013) observed a slight decrease in volume between 8 and 22 years. Discrepancies in findings across studies may arise due to differences in segmentation methods (e.g., manual versus automated), analytic approach (e.g., correcting versus not correcting for individual differences in brain volumes), or structure of the study (e.g., longitudinal versus cross-sectional).

The hippocampus is not a uniform structure. Notable differences in anatomical connectivity and genetic, molecular, and functional attributions have been identified along its longitudinal axis (Fanselow & Dong, 2010; Poppenk, Evensmoen, Moscovitch, & Nadel, 2013; Strange, Witter, Lein, & Moser, 2014; Vogel et al., 2020). Further, the hippocampal formation is comprised of histologically distinct subfields thought to make unique functional contributions to episodic memory (Marr, 1971). Accordingly, more recent studies have complimented their examination of whole hippocampal development by segmenting the hippocampus into sub-regions along the long axis (i.e., head, body, and tail or by anterior and posterior hippocampus; see (Canada et al., 2020; Canada et al., 2021; DeMaster et al., 2013; Joshua K Lee et al., 2015; Riggins et al., 2015) or subfields. Riggins and colleagues (2015) showed that 6-year-olds had marginally larger overall hippocampal volume than 4-year-old-children, but in a follow-up longitudinal study with 4- to 8-year-olds showed a quadratic pattern of change in the head of the hippocampus, with a more monotonic pattern in the body and tail (Canada et al., 2020). As noted earlier, Gogtay and colleagues (2006) did not observe age-related changes in overall hippocampal volume, but when examined along the long axis (i.e., anterior/posterior hippocampus) the anterior hippocampus showed an increase in volume with age, while the posterior hippocampus decreased in volume with age between 4 and 25 years.

Changes in hippocampal subfield volume have been identified across a broad range of ages. When examined from early-to-mid-childhood, the CA1 increased in volume between 4 and 5 years while the CA2-4/DG and subiculum peaked in volume between 5 and 6 years (Canada et al., 2021). Across a broader age range all studies have identified non-linear positive associations with age across hippocampal subfield peaking in adolescence (Krogsrud et al., 2014; Tamnes et al., 2018; Lee et al., 2014). Similarly, we observed non-linear increases in CA1 and DG/CA3 volumes with age that peaked in mid-teen years, but this pattern was not observed for the subiculum in our sample. Our results, taken together with prior studies, suggest that segmentation of the hippocampus into subfields or regions along its long axis provides a more accurate picture of developmental changes that can be masked when the hippocampus is examined as a whole.

Our findings add to the body of work suggesting a protracted hippocampal development of the hippocampus and subfields (with the exception of subiculum) at least into adolescence (Goddings et al., 2014; Joshua K Lee et al., 2015; Østby et al., 2009; Riggins et al., 2018; Tamnes et al., 2013; Uematsu et al., 2012; Yurgelun-Todd et al., 2003) While some studies have suggested that there is an early peak in hippocampal development or found no age related differences (Barnea-Goraly et al., 2014; Giedd et al., 1996; Gogtay et al., 2006; Joshua K Lee et al., 2015; Riggins et al., 2015; Uematsu et al., 2012) the differences in findings is potentially related to the variety of different age ranges studied. However, when more extended age ranges are included, hippocampal development seems to extend into mid-adolescence. Indeed, one shortcoming of previous research that the present study alleviates is the examination of varying ages within truncated time frames (DeMaster et al., 2013; Stine K Krogsrud et al., 2014; Joshua K Lee et al., 2014; Tamnes, Bos, van de Kamp, Peters, & Crone, 2018). The major contribution of the present study is the examination of a wide age range in a large sample with a good representation at each age range. This allows us to zoom out and take a “birds-eye view” of the development of the hippocampus, beginning from a very young age (3-years) through the adolescent years and into early adulthood.

Interestingly, even with our large sample we did not find that the entorhinal cortex followed the same trajectory of volumetric change over age as the hippocampus and its subregions. From one perspective, this may be surprising, given the extensive connectivity of the entorhinal cortex to the rest of the hippocampus. However, the results also suggest that volumetric changes can occur along differential timelines, even for highly connected regions. This is consistent with data from Alzheimer’s studies showing that changes in entorhinal cortex volume as a result of the disease do not immediately track with changes in hippocampal volume (Killiany et al., 2002). In terms of typical development, it is possible that pruning of hippocampal synapses follows a different timeline than pruning of neocortical synapses. Significant work has investigated axonal and dendritic pruning in the rodent hippocampus (Faulkner, Low, & Cheng, 2007). This work is far more difficult to investigate in the human hippocampus with histologic data, and thus changes in MRI volume must be used. However, even here we must be cautious. A recent paper has suggested that changes in MRI measurements over development (in this case, cortical thinning) may be more associated with changes in myelination, and not changes in grey matter morphometry, as was previously assumed (Natu et al., 2019). We speculate that it is possible that myelination is proceeding differently in the hippocampus than it is in the cortex, which could drive changes in volume as measured by MRI. However, further research would need to be conducted to support this claim. Thus, these two possible explanations may explain why we see developmental differences in the volume of hippocampus compared with the neighboring entorhinal cortex.

### 2. Hippocampal development in relation to the cortical grey matter

Hippocampal development progresses in the context of broader maturational changes across the brain. In the interest of investigating the relation between the hippocampal and cortical developmental trajectories, we examined changes in the *ratio* of hippocampal volume to whole brain cortical grey matter volume with age. If hippocampal development was following in lock step with the developmental course of the cortex than no discernable pattern would be evident across age. The linear pattern that emerged when examining the hippocampal-cortical ratio, on the contrary, highlights a tight relation between hippocampal and cortical development. Research on changes in whole brain cortical grey matter volume has demonstrated that cortical volume peaks earlier in childhood, and decreases in late childhood and throughout adolescence (Lebel & Beaulieu, 2011; Tamnes et al., 2013; Wierenga et al., 2014). The findings of the current study suggest that the hippocampus follows a somewhat inverse trajectory. This a first step however and the issue of how the hippocampus develops in relation to the broader cortical network to which it is connected deserves additional scrutiny.

### 3. Hippocampal volumetric associations with behavior

The hippocampus is an important region for episodic memory; thus, development of this structure should be related to memory performance in tasks that tax functions related to episodic memory. We found that bilateral hippocampus, as well as subfield volumes (CA1 and DG/CA3) were all positively associated with performance on the PSMT task, even after controlling for a number of relevant covariates (age, gender, ethnicity, parent highest level of education, household income, scanner type, and ICV). However, not all subfields exhibited a similar relation between volume and PSMT performance. It may be that specific subfields of the hippocampus play differential roles in the mechanics of episodic memory (Kim & Yassa, 2013; Wendelken et al., 2014), and this may differ across development (J. K. Lee et al., 2020). For example, research in patients with damage to the CA1 shows that they have extensive episodic memory loss suggesting an important role for CA1 in episodic memory (Zola-Morgan, Squire, & Amaral, 1986). In children, CA3/DG volume has been positively related to episodic memory (Lee et al., 2014) and greater volume in CA1 and CA2-3 has been related to recall and retention after an extended delay (Krogsrud et al., (2014). Our results are somewhat consistent with these previous studies. While we found that both CA1 and DG/CA3 volume (the latter bilaterally), predicted PTSM scores, we did not find the same relations for subiculum.

For the first time we show here that the ratio of the hippocampus to the rest of the cortex was positively associated with PTSM scores. Considered in the context of increasing hippocampal volume and concomitant cortical volume decrease with age, this finding suggests that children who retain or increase hippocampal volume as cortical volume declines exhibit better episodic memory. It remains to be seen whether the association between behavior and hippocampal development in the context of cortical maturation is specific to retention of hippocampal volume in the context of developmentally appropriate cortical reduction, increase of hippocampal volume in relation to steady cortical volume across age, or more likely a manifestation of the combination of hippocampal increases and cortical reductions across development – with those individuals exhibiting the steepest divergence between these two measures reaping the largest mnemonic enhancement.

### 4. White matter connectivity with the rest of the brain

In our final analysis we examined how several white matter pathways, which have been identified as playing important roles in episodic memory, contribute to episodic memory development. Unexpectedly, we did not find any evidence that these pathways are associated with developing episodic memory. A number of studies (Krogsrud et al., 2016; Loenneker et al., 2011; Moon et al., 2011; Simmonds, Hallquist, Asato, & Luna, 2014) have shown a protracted timeline of white matter development from early childhood until adulthood, with different rates of maturation in different pathways (Lebel, Treit, & Beaulieu, 2017 for review). We expected that such changes would be associated with behavioral changes in episodic development in hippocampus connectivity. Indeed, this has been found in some other studies (Mabbott et al., 2009; Ngo et al., 2017). For example, Ngo and colleagues (2017) examined the fornix and uncinate fasciculus in a small sample of 4- and 6-year-olds. Their results revealed that white matter pathways connecting hippocampus and inferior parietal lobule significantly predicted episodic memory performance. These data suggest that white matter connecting hippocampus and lateral regions is relevant to the development of episodic memory. Surprisingly, the current data are not consistent with these studies. One possible explanation for this may be that prior studies did not control for the covariates that we controlled for, including whole brain FA. This may have masked associations that were found in previous studies. However, we argue that such controls are necessary. For example, if whole brain white matter development is largely explanatory, it does not tell us anything about the specific fiber pathways connected to the hippocampus. In addition, studies based on small samples may actually inflate the reporting of effect sizes, especially in studies addressing brain-behavior associations (Dick et al., 2021). It is therefore possible that such relations reported in smaller samples are not reliable when larger samples are investigated. At the same time, this null finding was surprising and will need to be examined further in independent samples.

### 5. Limitations

Other methodological differences can also lead to different results, especially when it comes to examination of subfields of the hippocampus. The most ideal method of subfield segmentation is manual segmentation. Although this method has a lot of strengths, it is not feasible for examination of large publicly available datasets. Giuliano (2017) recently investigated this issue and showed that, indeed, different volumetric estimates are reported when using different methods, but the authors also showed that automated methods can give excellent results. The method we employed allowed us to exploit a data set with a larger sample size than would have been feasible using manual segmentation methods, but it is important to acknowledge this limitation. Thus, while we report differences with some prior research, the method we used is arguably appropriate for large-sample investigations, and indeed it is also consistent with a number of prior studies.

### 6. Summary

In summary, we found in a large sample of 830 3- to 21-year-olds, non-linear volume increases in the hippocampus and related subfields that peaked in the mid-teen years. These trajectories controlled for changes in the general size of the brain that also occur during childhood and adolescence. We also found significant age-related differences in global hippocampal volume when considered in the context of cortical grey matter volume. That is, we found that as cortical volume decreased with age, hippocampus volume continued to increase. In addition to age-related differences in grey matter volume, we found that several distinct subfields, specifically CA1 and DG/CA3, are significantly related to episodic memory as assessed using the PSMT. Finally, we examined how white matter pathways were associated with episodic memory, but did not find any evidence in our large sample to suggest that these pathways significantly influenced episodic memory development. These findings contribute to further understanding of hippocampal development over an extended age range, and its associated effects on episodic memory development.

